# HIV-PULSE: A long-read sequencing assay for high-throughput near full-length HIV-1 proviral genome characterization

**DOI:** 10.1101/2023.01.18.524396

**Authors:** Laurens Lambrechts, Noah Bonine, Rita Verstraeten, Marion Pardons, Ytse Noppe, Sofie Rutsaert, Filip Van Nieuwerburgh, Wim Van Criekinge, Basiel Cole, Linos Vandekerckhove

**Affiliations:** HIV Cure Research Center, Department of Internal Medicine and Pediatrics, Ghent University Hospital, Ghent University, 9000 Ghent, Belgium; BioBix, Department of Data Analysis and Mathematical Modelling, Faculty of Bioscience Engineering, Ghent University, 9000 Ghent, Belgium; Laboratory of Pharmaceutical Biotechnology, Faculty of Pharmaceutical Sciences, Ghent University, 9000 Ghent, Belgium

**Author notes:** Correspondence and requests for materials should be addressed to L.V. These authors contributed equally (Basiel Cole, Linos Vandekerckhove).

## Abstract

A deep understanding of the composition of the HIV-1 reservoir is necessary for the development of targeted therapies and the evaluation of curative efforts. However, current near full-length (NFL) HIV-1 proviral genome sequencing assays are based on labor-intensive and costly principles of repeated PCRs at limiting dilution, restricting their scalability. To address this, we developed a high-throughput, long-read sequencing assay called HIV-PULSE (HIV <u>P</u>roviral <u>U</u>MI-mediated <u>L</u>ong-read <u>Se</u>quencing). This assay uses unique molecular identifiers (UMIs) to tag individual HIV-1 genomes, allowing for the omission of the limiting dilution step and enabling long-range PCR amplification of many NFL genomes in a single PCR reaction, while simultaneously overcoming poor single-read accuracy. We optimized the assay using HIV-infected cell lines and then applied it to blood samples from 18 individuals living with HIV on antiretroviral therapy, yielding a total of 1,308 distinct HIV-1 genomes. Benchmarking against the widely applied Full-Length Individual Proviral Sequencing assay revealed similar sensitivity (11% vs 18%) and overall good concordance, though at a significantly higher throughput. In conclusion, HIV-PULSE is a cost-efficient and scalable assay that allows for the characterization of the HIV-1 proviral landscape, making it an attractive method to study the HIV-1 reservoir composition and dynamics.

## Introduction

The establishment of a viral reservoir shortly after HIV-1 infection leads to long-term viral persistence in people living with HIV-1 (PLWH) (1–3). While antiretroviral therapy (ART) can successfully suppress viral replication, it is not curative as the viral reservoir is not targeted (4, 5). Consequently, lifelong adherence to ART is required to prevent viral rebound, which usually takes place within several weeks following ART cessation (6). Despite the relatively low frequency of infected CD4 T cells that remain during ART (1/1,000 - 1/10,000), the size of the viral reservoir is remarkably stable, with an estimated half-life of 44 months (7, 8). The search for curative interventions targeting the viral reservoir remains one of the top priorities for achieving HIV-1 remission (9), however, this search is faced with two major challenges: (i) A lack of knowledge of the mechanisms governing HIV-1 latency and reservoir maintenance; (ii) A lack of high-throughput methods to measure the efficacy of reservoir-reducing interventions. To address these problems, technological advances that allow for a deep and high-throughput reservoir characterization are urgently needed (10).

Historically, the qualitative assessment of the HIV-1 reservoir has been carried out using two main types of assays: (i) Viral outgrowth assays (VOA), in which replication-competent viruses are reactivated and propagated ex vivo at limiting dilution, followed by quantification and sequencing of the cultured viral genomes (11, 12); (ii) Sequencing-based assays, where single proviral genomes are PCR-amplified from bulk DNA at limiting dilution, followed by Sanger- or short-read next-generation sequencing (NGS) (13–19). The VOA-based methods have the inherent benefit that they focus on replication-competent viruses, though they are usually long, costly and labor-intensive and have been shown to underestimate the true size of the replication-competent fraction following one round of reactivation (15). Sequencing-based methods allow the assessment of a small subgenomic region of interest or the near full-length (NFL) proviral genome (∼90%) (13–19). In the case of the latter, the percentage of genome-intact proviruses can be derived, which has previously been estimated at 2-5% on average (16). More recently, several flow-cytometry-based assays have been developed to isolate and study HIV-infected cells harboring an inducible provirus such as Simultaneous TCR, Integration site and Provirus sequencing (STIP-Seq), which specifically targets the translation-competent reservoir (20–22). In these assays, the infected cells are dispensed into single wells of a microtiter plate, followed by genomic or transcriptomic sequencing.

A common denominator of the assays described above is that they all rely on the physical isolation of individual viral genomes into different wells of a microtiter plate, followed by the PCR amplification of each genome in separate reactions (23). This principle is both labor-intensive and costly, severely limiting the applicability in large scale projects.

Up until the advent of long-read sequencing technologies, long amplicons (>5 kb) were either sequenced by a series of overlapping Sanger sequencing reactions, or by fragmentation of the amplicon into smaller pieces followed by short-read NGS (16–18). Long-read sequencing technologies offer the ability to sequence long amplicons in a single read, however, these technologies suffer from a high error rate of single-pass reads (∼5-10%) (24). Recently, Karst et al. developed a protocol that uses unique molecular identifiers (UMIs) to obtain high-accuracy consensus sequences from long amplicons (>5 kb), overcoming the problem of the limited single-read accuracy (25).

Here, we present a new assay that allows for high-throughput amplicon sequencing of NFL HIV-1 genomes, called HIV-PULSE: HIV <u>P</u>roviral <u>U</u>MI-mediated <u>L</u>ong-read <u>Se</u>quencing. By tagging individual HIV-1 genomes with two distinct UMIs, the step of limiting dilution can be omitted, enabling the amplification of many NFL genomes in a single reaction (25). After optimization of the assay on HIV-infected cell lines, we used the protocol to characterize the viral reservoirs of a chronic cohort of PLWH on ART (n=18). Benchmarking against the widely used Full-Length Individual Proviral Sequencing (FLIPS) assay revealed comparable accuracy and efficiency, but a remarkably higher throughput and lower cost per sequenced NFL HIV-1 genome (17). In conclusion, HIV-PULSE is a valuable addition to the arsenal of HIV-1 proviral sequencing methods and opens new possibilities for investigating the composition and dynamics of the HIV-1 reservoir.

## Materials and Methods

### Biological Resources

#### Study participants and sample collection

A total of 18 individuals on suppressive ART were included in this study (Supplemental Table 1). Participants were recruited at Ghent University Hospital and donated blood samples. Some participants agreed to additional leukapheresis to harvest large amounts of leukocytes. Peripheral blood mononuclear cells (PBMCs) were isolated by Ficoll density gradient centrifugation and were cryopreserved in liquid nitrogen. CD4 T cells were isolated from PBMCs by negative selection using EasySep Human CD4 T Cell Enrichment Kit (StemCell Technology, #19052). All participants signed informed consent forms approved by the Ethics Committee of the Ghent University Hospital (Belgium) (Ethics Committee Registration number: 2015/0894, 2016/0457, BC-07056).

#### Cell lines

J-Lat 8.4 cells (ARP-9847, contributed by dr. Eric Verdin), a Jurkat-based cell line latently infected with HIV, and Jurkat E6.1 cells (ARP-177, contributed by dr. Arthur Weiss) were obtained through the NIH HIV Reagent Program, Division of AIDS, NIAID, NIH (26, 27). J-Lat and Jurkat cells were grown in RPMI1640 (Gibco, #11875093) supplemented with 10% fetal bovine serum (HyClone, #RB35947) and 1% Pen/Strep (Gibco, #15140122).

#### DNA isolation and HIV-1 DNA reservoir size measurement

Genomic DNA from pelleted cells was isolated via column extraction using the DNeasy Blood & Tissue Kit (Qiagen, #69506) according to the manufacturer’s instructions. The DNA concentration of each extract was measured on a Qubit 3.0 fluorometer using the Qubit dsDNA BR assay kit (ThermoFisher Scientific, #Q32853). The HIV-1 copy number was determined by a total HIV-1 DNA assay on droplet digital PCR (Bio-Rad, QX200 system), as described previously (28). PCR amplification was carried out with the following cycling program: 10 m at 95°C; 40 cycles (30 s at 95°C, 1 m at 56°C); 10 m at 98°C. Droplets were read on a QX200 droplet reader (Bio-Rad). Analysis was performed using ddpcRquant software (29).

#### Full-length individual proviral sequencing

The full-length individual proviral sequencing (FLIPS) assay was performed as described by Hiener et al. (17). A detailed protocol can be found on the following link: dx.doi.org/10.3791/58016. In short, genomic DNA was used as input for a nested PCR performed at an endpoint dilution where <30% of the reactions are positive. The cycling conditions were 94°C for 2 m; then 94°C for 30 s, 64°C for 30 s, 68°C for 10 m for 3 cycles; 94°C for 30 s, 61°C for 30 s, 68°C for 10 m for 3 cycles; 94°C for 30 s, 58°C for 30 s, 68°C for 10 m for 3 cycles; 94°C for 30 s, 55°C for 30 s, 68°C for 10 m for 21 cycles; then 68°C for 10 m. For the second round, 10 extra cycles at 55°C were included. The PCR products were visualized using agarose gel electrophoresis (1% agarose gel). Proviral amplicons from positive wells were cleaned using AMPure XP beads (Beckman Coulter, #A63880), followed by a quantification of each cleaned provirus with Quant-iT PicoGreen dsDNA Assay Kit (Invitrogen, #P11496).

#### Illumina short-read sequencing

Selected FLIPS proviral amplicons underwent NGS library preparation using the Nextera XT DNA Library Preparation Kit (Illumina, #FC-131-1096) with indexing of 96-samples per run according to the manufacturer’s instructions (Illumina, #FC-131-2001), except that input and reagents volumes were halved and libraries were normalized manually. The pooled library was sequenced on a MiSeq Illumina platform via 2×150 nt paired-end sequencing using the 300 cycle v2 kit (Illumina, #MS-102-2002).

#### HIV-PULSE assay methodology

##### Pre-amplification

A first PCR was used to specifically target and pre-amplify HIV-1 proviral templates using the outer primers of a nested HIV-1 primer set (listed in Supplemental Table 2, Figure 1A). Each PCR reaction contained 500 ng of genomic DNA, 2 µL LongAmp Taq DNA Polymerase (NEB, # M0323L), 0.5 µM of each primer (First PCR F, First PCR R), 1.5 µL 10 mM dNTPs (Promega, #C1141), 10 µL 5X LongAmp Taq Reaction Buffer in 50 µL. The following cycling conditions were used: 94°C for 1 m 15 s; then 94°C for 30 s, 63°C for 30 s, 65°C for 10 m for 6 cycles; then 65°C for 10 m. The number of amplification cycles can be reduced to 5 cycles for samples from individuals with a high reservoir size (>2,500 total HIV-1 DNA copies/million CD4) to prevent overbinning. PCR products were cleaned using CleanPCR magnetic beads (CleanNA, #CPCR-0050) at a 1.0x beads/sample ratio.

**Figure 1.**
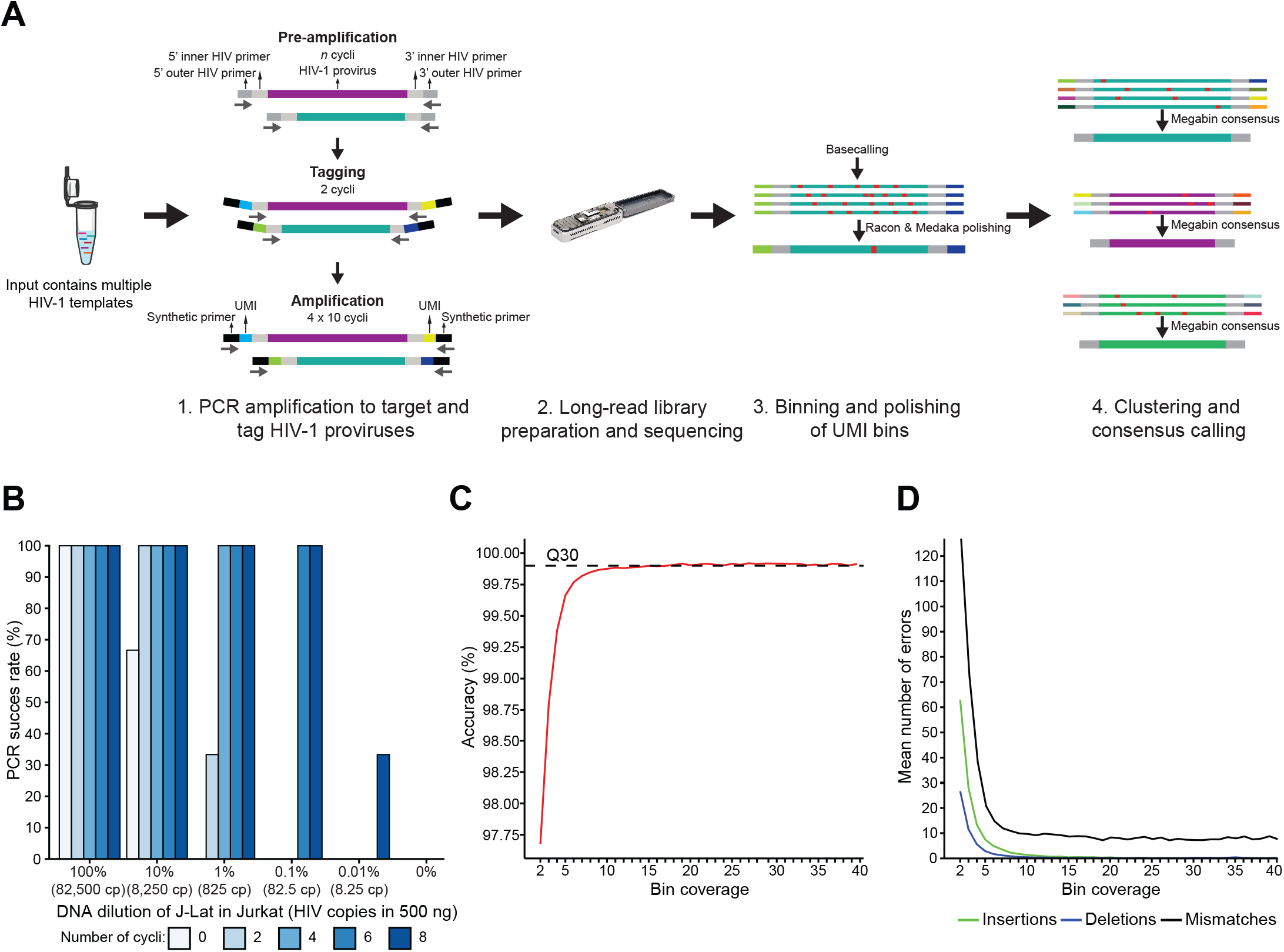
HIV-PULSE methodology overview and performance evaluation. (A) Schematic overview of the HIV-PULSE assay. A PCR reaction with bulk DNA containing multiple HIV-1 templates is pre-amplified using outer HIV-1 primers for a limited number of cycles to improve sensitivity. Next, pre-amplified material is tagged with a dual barcode consisting of a unique molecular identifier (UMI) attached to both ends using an HIV-1 specific inner primer. To generate enough material for long-read library preparation, the tagged material is amplified with synthetic primers in several PCR rounds followed by clean up to prevent length bias. (B) Success rate of the HIV-PULSE assay for different input ratios of HIV-1 with varying number of PCR cycles during pre-amplification. Each condition was performed in triplicate. (C) Mean accuracy of HIV-PULSE bin consensus sequences with increasing bin coverage compared to the Illumina reference sequence. The dashed line indicates the Q30 (99.9% accuracy) threshold. (D) Mean number of errors (insertions, deletions and mismatches) found in HIV-PULSE consensus sequences of 9.5 kb with increasing bin coverage compared to the Illumina reference sequence.

##### Tagging HIV-1 templates

A second PCR was performed to tag both ends of the pre-amplified proviral HIV-1 templates with a tailed UMI (listed in Supplemental Table 2). Primers were designed to contain: (i) a synthetic primer binding site used in later stages for amplification, (ii) a UMI with a repetitive pattern of 12 random nucleotides and 6 degenerate nucleotides (Y/R) and (iii) an HIV-1 inner primer of the nested primer set to target the pre-amplified templates (Figure 1A, Supplemental Figure 1A). Each PCR reaction contained all the cleaned pre-amplified product (30 µL), 2 µL LongAmp Taq DNA Polymerase (NEB, # M0323L), 0.5 µM of each primer (Second PCR F UMI, Second PCR R UMI), 1.5 µL 10 mM dNTPs (Promega, #C1141), 10 µL 5X LongAmp Taq Reaction Buffer in 50 µL. The following cycling conditions were used: 94°C for 1 m 15 s; then 94°C for 30 s, 58°C for 30 s, 65°C for 10 m for 2 cycles; then 65°C for 10 m. Tagged PCR products were cleaned using CleanPCR magnetic beads (CleanNA, #CPCR-0050) in a custom buffer solution (based on the ‘SPRI size selection protocol for >1.5–2 kb DNA fragments’ protocol provided by Oxford Nanopore Technologies (ONT)) at a 0.9x beads/sample ratio and eluted in 30 µL nuclease-free water (NFW).

##### Amplification of UMI-tagged proviruses

The next steps used 4 consecutive PCR amplification rounds of each 10 cycles followed by a clean up to produce enough template input required for long-read sequencing while preserving amplicon size distributions. Here, we made use of a primer set that binds to the synthetic binding site incorporated during the previous tagging stage. The PCR mix consists of 2 µL LongAmp Taq DNA Polymerase (NEB, # M0323L), 0.5 µM of each primer (ncec_pcr_fw_v7, ncec_pcr_rv_v7), 1.5 µL 10 mM dNTPs (Promega, #C1141), 10 µL 5X LongAmp Taq Reaction Buffer in 50 µL. For the first PCR amplification round all the cleaned tagging products from the previous step (30 µL) were used as template input while during the second, third and fourth amplification rounds only a third of the cleaned product of the previous round is used (10 µL). The following cycling conditions were used: 94°C for 1 m 15 s; then 94°C for 30 s, 58°C for 30 s, 65°C for 10 m for 2 cycles; then 65°C for 10 m. PCR products were cleaned after each consecutive round using regular CleanPCR magnetic beads (CleanNA, #CPCR-0050) at a 1.0x beads/sample ratio and eluted in 30 µL NFW. During the last round of 10 cycles, the regular primers were switched for a custom set of tailed primers to barcode the PCR products from the same participant with a specific, identical identifier (listed in Supplemental Table S2). After the last PCR round, the end products were visualized using agarose gel electrophoresis (1% agarose gel) and the DNA concentration was determined using a Qubit 3.0 fluorometer with the Qubit dsDNA BR assay kit (ThermoFisher Scientific, #Q32853).

### ONT long-read sequencing

Samples were multiplexed using the Native Barcoding Kit (ONT, #EXP-NBD104) using the following strategy: each PCR replicate was assigned to a different ONT barcode and contained equimolarly pooled PCR products from different participants. In later stages, this allows to assign reads to the correct PCR replicate by the ligated ONT barcode and to the correct sample by the participant-specific identifier attached during the last PCR round (Supplemental Figure 1A). For library preparation, the Ligation Sequencing Kit (ONT, #SQK-LSK109) was used following the manufacturer’s instructions. Samples were sequenced on a MinION ONT device using MinION R10.3 flow cells and the MinKNOW v21.02.1 software followed by basecalling at super accuracy mode and demultiplexing with Guppy v5.0.17.

### Bioinformatics analysis of long-read data

For the analysis of long-read data, a customized version of the UMI data analysis workflow as described by Karst et al. was utilized (Supplemental Figure 1B) (25). The adapted scripts can be found at https://github.com/laulambr/longread_umi_hiv, main changes include updated software versions of samtools (1.11), medaka (1.4.3) and racon (1.4.20). Before the data was analyzed using the UMI pipeline, the demultiplexed ONT reads were first mapped against the HXB2 reference sequence using minimap2 (2.17) to filter out non-HIV-1 reads. Next, the ‘*longread_umi nanopore_pipeline*’ workflow was run separately on each replicate read dataset using the following settings: *-s 200 -e 200 -m 1500 -M 10000 -f CAAGCAGAAGACGGCATACGAGAT -F AAGTAGTGTGTGCCCGTCTGTTGTGTGAC -r AATGATACGGCGACCACCGAGATC -R GGAAAGTCCCCAGCGGAAAGTCCCTTGTAG -c 3 -p 1 -q r103_hac_g507 -U ‘r103_min_high_g360’*. The workflow consists of the following consecutive steps; (i) trimming and filtering of the HIV-1 long-read sequencing data using Porechop (v.0.2.4, https://github.com/rrwick/Porechop), Filtlong (v.0.2.0, https://github.com/rrwick/Filtlong) and cutadapt (v.2.7) (30), (ii) extraction of UMI reference sequences using cutadapt (v.2.7) and usearch (30, 31), (iii) binning of reads to UMI combinations using bwa (v0.7.17) and samtools (v1.11) while excluding chimeric artifacts, (iv) generation of bin consensus sequences using usearch and minimap2 (v2.17) (31, 32) and (v) polishing of bin consensus data by multiple rounds of racon (v.1.4.20) and a final round of Medaka (v.1.4.3, https://github.com/nanoporetech/medaka) (33).

Next, a custom bioinformatics workflow specific to the HIV-PULSE protocol was run to correct for pre-amplification, improve final consensus accuracy and evaluate clonality among PCR replicates. First, the polished bin consensus sequences from each replicate dataset were demultiplexed to their respective participant using the ‘*longread_umi demultiplex*’ workflow. For each participant, bin consensus sequences from all PCR replicates were pooled together and screened for the occurrence of identical UMIs. Identical UMI pairs in different PCR replicates are technically not possible but these artifacts may arise due to misassignment during the initial raw ONT reads demultiplexing by Guppy. In these cases, the bin in the replicate with the highest coverage was considered correct while the others were removed from their respective replicate datasets. As the assay relies on a pre-amplification phase, single proviral templates will have been amplified and potentially tagged into bins with different UMI pairs. This prohibits the screening for clonal proviruses present in one bulk PCR reaction as pre-amplification would also lead to the presence of identical proviruses in bins with different UMI pairs. To correct for this, identical proviruses (due to clonality or pre-amplification) present in the bin consensus sequences from each participant were identified by clustering genomes with similar sizes and >99.5 % sequence identity into a megabin using usearch (31). For megabins consisting of >3 bins, a new consensus sequence was constructed to help to resolve remaining errors. For megabins that only consisted of 2 bin consensus sequences, the bin with the highest coverage (∼accuracy) was retained while bins that did not cluster remained as unique bins. Finally, an assessment of clonality among different PCR replicates was made by screening the resulting megabins for the presence of bin sequences originating from different PCR replicates. If the same proviral sequence was found in multiple PCR replicates, it is considered as evidence of a potential clonal origin, as opposed to proviruses that are only found in one replicate. By performing multiple PCR replicates, one can identify clonal populations, however, accurate quantification of reservoir clonality is hindered by the fact that identical proviruses found within the same PCR replicate are collapsed and counted as one (to exclude potential bias by pre-amplification).

### Bioinformatics analysis of Illumina data

NFL genome sequences derived from FLIPS reactions were de novo assembled as follows: (i) FASTQ quality checks were performed with FastQC v0.11.7 (http://www.bioinformatics.babraham.ac.uk/projects/fastqc) and removal of Illumina adaptor sequences and quality-trimming of 5′ and 3′ terminal ends was performed with bbmap v37.99 (https://sourceforge.net/projects/bbmap/). (ii) Trimmed reads were de novo assembled using MEGAHIT v1.2.9 with standard settings (34). (iii) Resulting contigs were aligned against the HXB2 HIV-1 reference genome using blastn v2.7.1 with standard settings, and contigs that matched HXB2 were retained (35). (iv) Trimmed reads were mapped against the *de novo* assembled HIV-1 contigs to generate final consensus sequences based on per-base majority consensus calling, using bbmap v37.99 (https://sourceforge.net/projects/bbmap/). Scripts concerning de novo assembly of HIV-1 genomes can be found at the following GitHub page: https://github.com/laulambr/virus_assembly.

### HIV-1 genome classification

NFL proviral genome classification was performed using the publicly available “Gene Cutter” and “Hypermut” webtools from the Los Alamos National Laboratory HIV sequence database (https://www.hiv.lanl.gov). Proviral genomes were classified in the following sequential order: (i) “Inversion”: presence of internal sequence inversion, defined as region of reverse complementarity. (ii) “Large internal deletion”: internal sequence deletion of >1000□bp. (iii) “Hypermutated”: APOBEC-3G/3F-induced hypermutation. (iv) “packaging signal and/or major splice donor (PSI/MSD) defect”: deletion >7□bp covering (part of) the packaging signal region or absence of GT dinucleotide at the MSD and GT dinucleotide at the cryptic donor site (located 4□bp downstream of MSD) (19). Proviruses with a deletion covering PSI/MSD that extended into the *gag* gene, thereby removing the *gag* AUG start codon, were also classified into this category. (v) “Premature stop-codon/frameshift”: premature stop-codon or frameshift caused by mutation and/or sequence insertion/deletion in the essential genes *gag, pol* or *env*. Proviruses with insertion/deletion >49 nt in *gag*, insertion/deletion >49□nt in *pol*, or insertion/deletion >99 nt in *env* were also classified into this category. (vi) “Intact”: proviruses that displayed none of the above defects were classified into this category.

### Reference sequences

Proviral HIV-1 genomes from participants previously acquired with the FLIPS and STIP-Seq assays in the context of former studies were included as references (20). Accuracy metrics on the new HIV-PULSE assay compared to FLIPS and STIP-Seq reference proviruses (sequenced using Illumina technology) were calculated using pomoxis (v0.3.6, https://github.com/nanoporetech/pomoxis).

### Phylogenetic analysis

Sequences obtained with the HIV-PULSE assay, STIP-Seq and FLIPS were multiple aligned using MAFFT (v 7.471) (36). DIVEIN was used to calculate pairwise diversity among proviral sequences (37). Phylogenetic trees were constructed using PhyML v3.0 (best of NNI and SPR rearrangements) and 1,000 bootstraps (38). R (v4.1.2), ggplot (v3.3.5) and ggtree (v3.2.1) were used for visualization and annotation of trees (39–41).

## Results

### Optimization of PCR cycling conditions on HIV-infected cell lines

Since the frequency of HIV-1 infected cells in ART-suppressed PLWH is remarkably low, typically in the range of 100-1,000 proviral genomes per million CD4 T cells, we set out to devise an assay allowing for the detection of such rare events using a UMI-binning strategy (Figure 1A) (7). By including a pre-amplification step, the sensitivity of the assay should considerably improve as more target templates are created, thus increasing the number of tagged templates transferred to the PCR amplification step. This hypothesis was tested by preparing a dilution series of J-Lat 8.4 DNA (HIV-infected CD4 T cell line) in Jurkat DNA (non-infected parental CD4 T cell line). The dilution series, ranging from 100% to 0.01% J-Lat 8.4 DNA (Jurkat as negative control), was subjected to either 0, 2, 4, 6 or 8 cycles of pre-amplification, using primers that target NFL HIV-1 (Figure 1B). Each amplification was performed in triplicate, using a fixed input of 500 ng genomic DNA (corresponding with ∼82,500 CD4 T cells) with the PCR success being determined by visualization of the ∼9.5 kb amplicon by agarose gel electrophoresis. As expected, reactions with an input of undiluted J-Lat DNA were all successful regardless of the number of pre-amplification cycles (Figure 1B). However, in samples with a proviral burden closer to those observed in samples from PLWH (∼0.1%) at least 6 cycles of pre-amplification were required for guaranteed PCR success.

Next, we set out to determine whether high-accuracy HIV-1 genomes could be acquired by performing long-read sequencing of the J-Lat 8.4 PCR products and subsequent analysis using a custom version of the UMI pipeline developed by Karst et al. (25). Using an ONT R10.3 flow cell, reads with a median length of 9.5 kb were generated, in agreement with the expected amplicon length of J-Lat 8.4 DNA (Supplemental Figure 1C-H). The bioinformatics pipeline allowed for the construction of the UMI-tagged proviruses by binning the reads based on the observed terminal UMI pairs in the sequencing data (Figure 1A). During these steps, non-HIV reads were filtered out (removing non-specific amplicons) and aberrant UMI bins not meeting a fixed list of criteria (*e*.*g*. chimeras, read orientation bias) were excluded (Supplemental Figure 1B-H, for LIB01 (sequencing library containing the J-Lat 8.4 amplicon reads) ∼30% of all detected UMI pairs did not meet the criteria, however, they accounted only for 16% of all the binned reads). For each bin, a consensus sequence was constructed by multiple rounds of polishing (racon, medaka) of the assigned raw reads. To assess the accuracy of the proviral UMI consensus sequences, we compared each bin to a reference genome of the J-Lat 8.4 amplicon sequenced with Illumina. As more reads were assigned to a bin, an increase in the mean accuracy could be observed before reaching a plateau at 99.9% (Figure 1C). Bins with a coverage of at least 15 reads passed the Q30 (99.9% accuracy) threshold. At the Q30 threshold, an average of 8 mismatches per 9.5 kb genome could be observed, while aberrant deletions and insertions were completely resolved (Figure 1D).

### Application of the assay on samples from PLWH and comparison to FLIPS

To assess the performance of the HIV-PULSE assay on samples from ART-suppressed PLWH, and to benchmark it to a gold standard short-read sequencing assay, DNA from peripheral blood CD4 T cells from 4 individuals was subjected to both the HIV-PULSE and the FLIPS assay (Figure 2). This yielded a median of 18 FLIPS and 87 HIV-PULSE distinct proviruses per participant (Figure 2A and Supplemental Figure 2). Of note, in the case of FLIPS, 10 out of 136 HIV-1 PCR amplicons failed to produce a correct HIV-1 genome during *de novo* assembly (Supplemental Figure 2A, median of 2.5 assembly failures per participant). Across both assays, a total of 16 overlapping expansions of identical sequences (EIS) were observed (Figure 2A). HIV-PULSE successfully identified 81% (13/16) of the overlapping EIS as being clonal, while FLIPS detected 37.5% of the EIS (6/16). However, this discrepancy is probably the result of the lower number of genomes assayed with FLIPS (∼5-fold). We compared the overlapping proviral sequences to estimate sequencing accuracy and link it to proviral classification. The mean accuracy was found to be 99.99% (Q40) for the megabinned proviruses, with residual errors observed in only four genomes due to homopolymeric regions (Supplemental Figures 2B-C). However, these errors did not affect the correct HIV-1 proviral classification.

**Figure 2.**
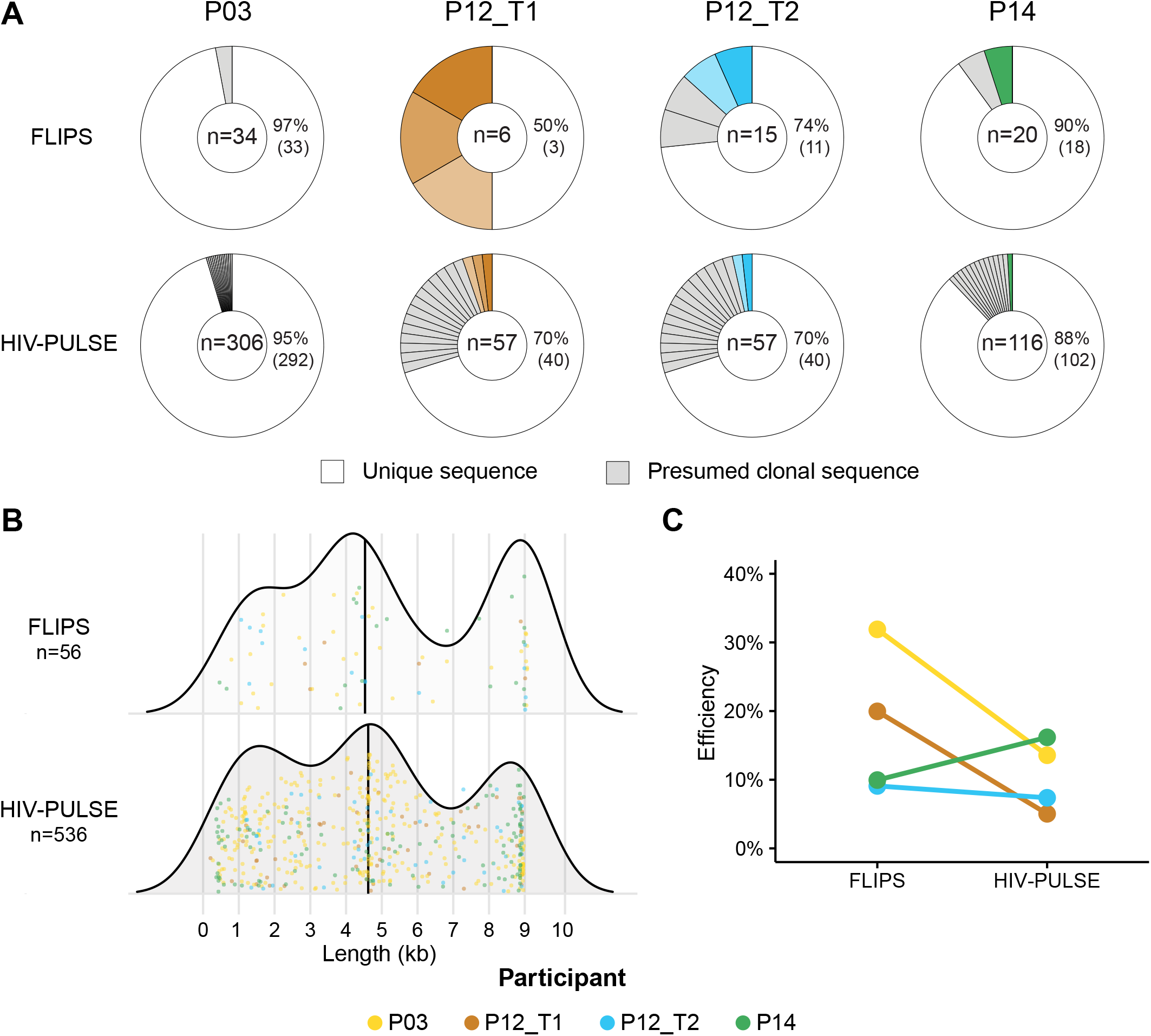
Benchmarking assays: novel HIV-PULSE vs gold standard FLIPS. (A) Donut plots displaying the fraction of unique and presumed clonal proviral sequences detected in each participant for both assays. The number of distinct proviruses generated by each assay is shown in the middle of each donut. The matching colored slices indicate the 6 out of 16 overlapping expansions of identical sequences (EIS) found to be clonal in both assays. p values test was used for a difference in the proportion of unique proviruses between both assays by ‘‘prop.test’’ in R, none were significant (*p*=1.00, *p*=0.583, *p*=1.00, *p*=1.00 for P03, P12_T1, P12_T2 and P14, respectively). (B) Size distributions of the observed proviral genome lengths for each assay. No significant difference was observed between both assays using a Kruskal-Wallis test (*p*= 0.08099). Each dot represents a single distinct provirus and is given a color for each participant. (C) For each assay and participant, the percentage of detected proviruses out of the total HIV-1 DNA reservoir size is shown. Assay efficiencies were compared for significance using a Kruskal-Wallis test -test (*p*=0.248).

Overall, comparing HIV-PULSE results to FLIPS data reveals (i) No significant differences between the proportions of sampled unique proviruses with each assay (Figure 2A; *p*=1.00, *p*=0.583, *p*=1.00, *p*=1.00 for P03, P12_T1, P12_T2 and P14, respectively) and (ii) No significant bias in the size distribution of the observed proviral genome lengths (Figure 2B, median lengths are 4,620 for HIV-PULSE and 4,531 for FLIPS, *p*=0.0810). The efficiency of both methods was assessed by calculating the percentage of total detected proviruses out of the total HIV-1 DNA reservoir size (Figure 2C). Slightly lower efficiencies were observed with the HIV-PULSE assay (mean of 11% opposed to 18% with FLIPS, *p*=0.271), however, this measure is likely an underestimation as it does not account for clonality of the reservoir (true size of clones is missed).

In conclusion, these results indicate that HIV-PULSE displays the required sensitivity for the amplification of NFL HIV-1 genomes from samples of ART-suppressed PLWH. Furthermore, both the accuracy and efficiency of the assay are on par with the FLIPS assay.

### HIV-PULSE enables high-throughput sequencing of NFL genomes in a large cohort of PLWH

The assay was next applied to peripheral blood CD4 T cell DNA of 18 chronically treated PLWH (mean time on ART = 11.2 years, Supplemental Table 1). The HIV-PULSE assay yielded an average of 15 ± 3 distinct HIV-1 proviruses per replicate, per participant (range: 3-55). For each participant, a mean PCR success rate of 97% was observed among the 6 PCR replicates based on agarose gel visualization (Supplemental Table 3). Overall, a total number of 1,661 proviruses (1,308 distinct) were retrieved across all participants (mean of 87 HIV-1 proviruses per sample, Supplemental Table 3, Supplemental Figure 3). Excluding the effect of clonal proliferation on infected cells, we looked at the presence of putatively intact genomes within the 1,308 distinct proviruses. A mean proportion of 5% intact distinct genomes was found across the 19 samples, which corresponds to previously reported numbers (Figure 3A) (16–18). Putatively intact sequences found across multiple replicates, indicative of clonality, were seen in 9/14 participants with at least 1 distinct intact sequence (Figure 3B). As two collected samples belonged to the same individual (P12) taken 3 years apart, a longitudinal assessment of the reservoir composition could be performed. This revealed the persistence of 21 infected cell clones (two with an intact provirus) between the two sampled timepoints (Figure 3, Supplemental Figure 4). No significant differences were observed in both the yield (*p*=0.662, 19 at T1 vs 18 at T2 mean distinct viruses per replicate, Figure 3A, Supplemental Table 2) and the observed mean pairwise distance among intact sequences between the two timepoints (P12_T1: 0.0230 n=7 vs P12_T2: 0.0197, n=5).

**Figure 3.**
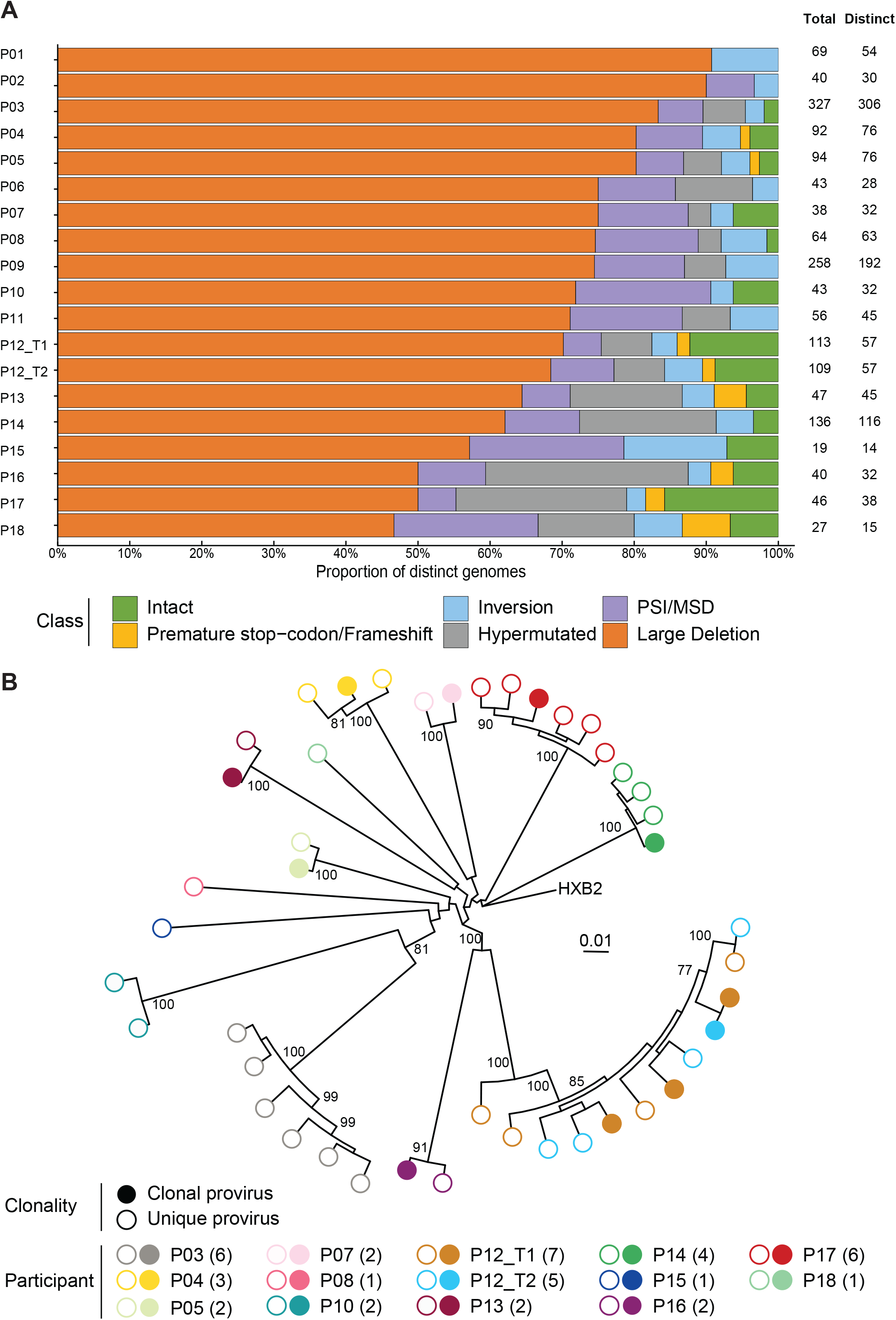
Proviral reservoir as assayed by the HIV-PULSE assay for a chronic cohort. (A) The proportions of different proviral classes observed among the distinct proviruses for each participant. On the right the number of total and distinct proviruses is displayed for each participant. (B) A phylogenetic tree including the distinct genome intact sequences. Each participant is shown as different colored dots, empty symbols indicate sequences only found once (unique, white insert) in a PCR replicate.

**Figure 4.**
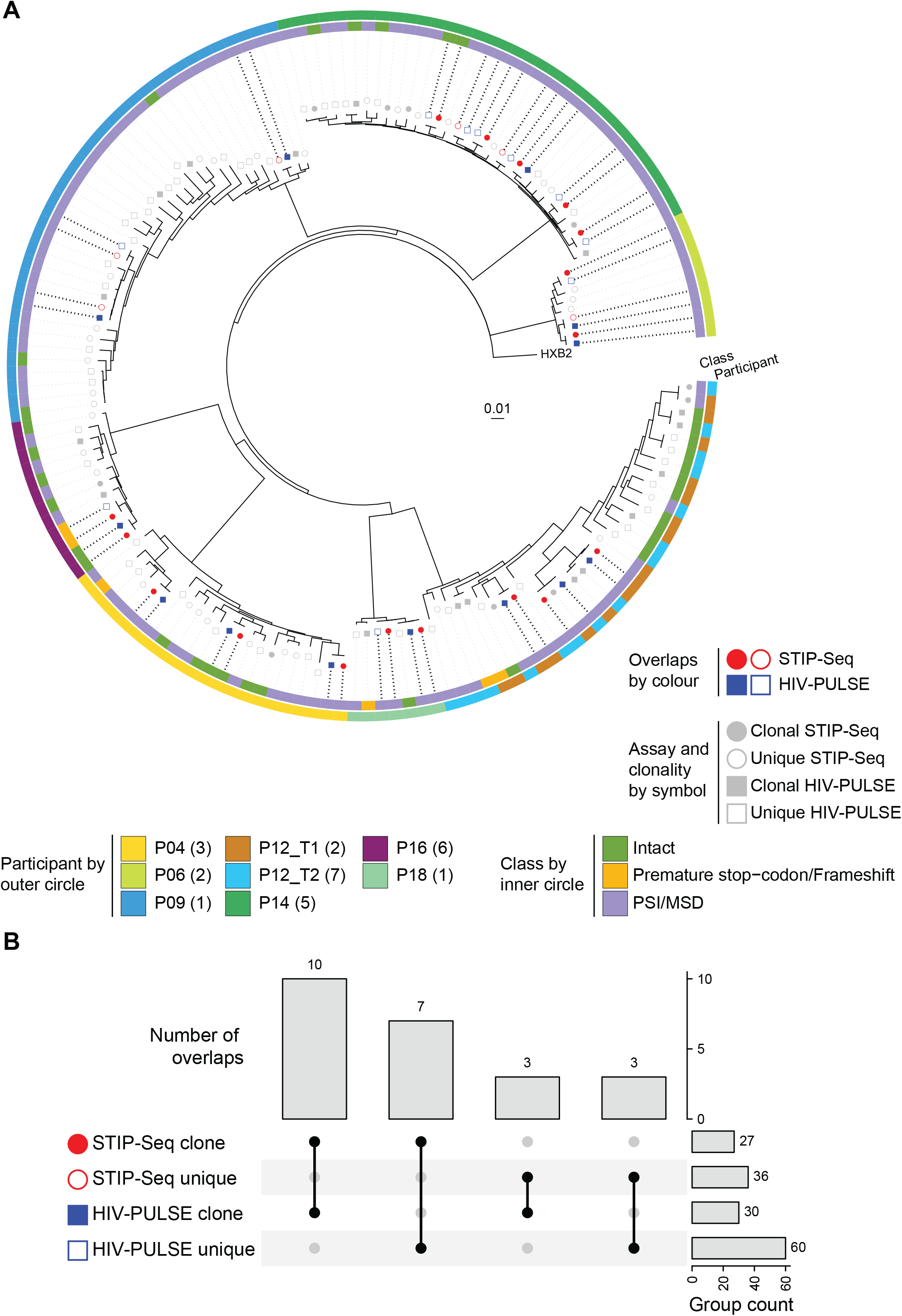
Benchmarking of HIV-PULSE vs STIP-Seq assay. (A) Phylogenetic tree including all distinct proviruses obtained with the HIV-PULSE (excluding sequences with inversions, large deletions and hypermutations) and STIP-Seq assays for 8 participants. Symbols reflect the different assays, proviruses only recovered in a single assay are shown in grey while assay overlapping are shown in red (STIP-Seq) or blue (HIV-PULSE). Empty symbols indicate sequences were found once (unique, white insert) in that respective assay. The outer and inner circles indicate for each provirus respectively the participant origin and associated HIV-1 genome classification. (B) UpSet-plot visualizing the number of overlaps between clonal and unique proviruses recovered with each respective assays.

While the characteristics of the cohort are quite diverse, we were still able to get a fair number of proviral sequences in individuals with a low total HIV-1 reservoir size (<500 total HIV DNA copies/million CD4, n=3). A significant correlation between the HIV-PULSE yield and the reservoir size measured by total HIV DNA was observed (Supplemental Figure 5, R=0.71, *p*=1.5*10^15^). On average, the efficiency of HIV-PULSE for these samples was 13% (95% CI [7.9, 17.6]) as calculated by dividing the number of detected distinct proviruses by the total number of input HIV-1 copies per PCR replicate (Supplemental Table 3). Nevertheless, this measure of efficiency is an underestimation, as it does not account for clones found within a replicate.

### HIV-PULSE detects proviruses of the translation-competent HIV-1 reservoir

While the proviral reservoir consists of a heterogeneous mix of HIV-1 proviruses belonging to different classes, only a few can contribute to viral rebound and/or HIV-1 pathogenesis by inducing chronic immune activation (*e*.*g*. intact, PSI/MSD defect) (20, 42, 43). Of particular clinical interest, the translation-competent HIV-1 reservoir represents all proviruses that can produce viral proteins following maximal stimulation, consequently enriching for replication-competent proviruses (42). In this regard, we next set out to evaluate whether HIV-PULSE can capture proviruses that belong to the translation-competent reservoir, by comparing HIV-PULSE data (n=8 participants) to STIP-Seq data (Figure 4A). On average per individual, 69% (95% CI [45, 92]) of the translation-competent STIP-Seq clones were detected with HIV-PULSE (as unique or clone). This corresponds to a total of 17 overlaps among both assays, of which 10/17 (59%) were also identified as a clone by HIV-PULSE, as they were detected across multiple replicates from the same participant (Figure 4B). Additionally, 17 clones were found in the HIV-PULSE data but not sampled with STIP-Seq (6/17 intact), which could be due to the integration of those proviruses at a chromosomal location in regions with features (*e*.*g*. heterochromatin) associated with HIV-1 transcriptional latency or a location less prone to be induced by latency reversing agents or by the more limited number of proviruses assessed with STIP-Seq compared to HIV-PULSE (44, 45). In conclusion, HIV-PULSE reliably picks up proviruses that belong to the translation-competent reservoir, which is of high importance for applicability in a clinical setting.

## Discussion

To achieve an HIV-1 cure, a comprehensive understanding of the persisting viral reservoir is crucial. Over the recent years, the application of HIV-1 NFL sequencing assays has increased our knowledge of certain key aspects, such as the proviral genomic composition and reservoir dynamics within PLWH. Still, all these results have been obtained through labor-intensive assays relying on limiting dilution and subsequent one-by-one sequencing of proviral genomes, limiting their application in large-scale studies. Here, we present the HIV-PULSE assay, allowing for a scalable, high-throughput assessment of the proviral HIV-1 reservoir. The use of dual barcodes removes the need for limiting dilution, allowing the amplification of multiple proviral HIV-1 templates during long-range NFL PCR on bulk DNA, while overcoming the inherently low single-read accuracy of long-read sequencing technologies. Benchmarking against the gold standard FLIPS method revealed comparable accuracy and efficiency, though notable differences in terms of throughput and associated costs are apparent. An overview of the approximated cost (in USD) per proviral sequence for each method indicates a 10-fold price reduction in favor of HIV-PULSE (Supplemental Table 4). Aside from a clear cost benefit, eliminating the limiting dilution step and sampling multiple proviruses out of a single PCR reaction offers a more high-throughput approach to NFL HIV-1 reservoir characterization. To illustrate, at least 52 96-well FLIPS PCR plates at limiting dilution would be required to obtain the equivalent of 1,661 total HIV-PULSE proviruses (Supplemental Table 4). Also, as a consequence of employing short-read NGS, FLIPS requires a *de novo* assembly step, which sometimes fails when resolving more complex genomes.

In this study, we applied HIV-PULSE to peripheral blood samples from a cohort of 18 PLWH on chronic ART, to study the composition of their HIV-1 viral reservoir. Out of the 1,308 distinct proviruses detected, ∼5% were deemed genome-intact, agreeing with earlier reports on PLWH on chronic ART (16). We compared the HIV-PULSE assay to the STIP-Seq assay for 8 participants, showing that HIV-PULSE efficiently picked up translation-competent proviruses, detecting 69% of the clonal sequences found with STIP-Seq. Interestingly, HIV-PULSE detected additional putatively intact proviral genomes which were not detected with the STIP-Seq assay, potentially representing hard-to-reactivate proviruses in a deep state of latency. While the HIV-PULSE assay does not enable a specific enrichment of the translation-competent reservoir, these findings make a case for HIV-PULSE as a tool to perform qualitative in-depth characterization of the functional reservoir dynamics in response to curative interventions during clinical trials. The high-throughput and cost-efficient nature of the HIV-PULSE assay makes it an attractive method for use within large-scale clinical studies of the HIV-1 reservoir with applications ranging from performing a longitudinal phylogenetic analysis of the proviral reservoir, screening multiple samples from the same individual for compartmentalization across different tissues, drug-resistance screening and bNAb epitope mapping. Indeed, implementing a qualitative NFL approach in clinical trial settings could help to check whether participants are eligible by excluding pre-existing resistance to a compound of interest or to assess immune escape following the intervention (46–48).

The adoption of long-read sequencing technologies for amplicon sequencing has historically been taken aback by the poor single-read accuracy. Notwithstanding these initial reservations, others have been developing long-read assays to characterize different aspects of the HIV-1 reservoir over the last couple of years. Pooled CRISPR Inverse PCR sequencing (PCIP-seq) allows to study both the integration site and the associated provirus using a targeted enrichment and inverse PCR strategy (49). While the collected data is certainly informative, the approach is hampered by limited sensitivity (3.2%), inadequate proviral coverage for accurate genome assembly and its reliance on the design of a custom pool of CRISPR guide RNAs for each participant. In comparison, HIV-PULSE has an improved sensitivity (13%), produces high-accuracy genomes, and does not rely on individualized primer designs, although information on the genomic location of the integrated provirus is missing. Another group developed NanoHIV, a bioinformatics tool to construct HIV-1 consensus sequences from long-read ONT data (50). This follows a reference mapping-based strategy with consecutive mappings to refine the original draft and deal with variable genomic regions. To compare the performance, they generated consensus genomes of NFL amplicons (acquired via nested NFL PCR performed at limiting dilution) and performed sequencing with both ONT and NGS Illumina. The authors report a mean accuracy of 99.4% (or 54 errors in a 9 kb genome), considerably lower than the megabin accuracy of 99.99% (or 1 error in a 9 kb genome) reported with our bioinformatics pipeline. Two studies describe protocols to amplify and sequence different genomic regions with accuracies up to 99.9% from virions in plasma samples from viremic individuals. While one relies on circular consensus sequencing (CSS) reads with PacBio technology to obtain 2.6 kb full-length *env* sequences (51), the other method Multi-read Hairpin Mediated Error-Correction Reaction (MrHAMER) targets a 4.6 kb *gag-pol* region followed by sequencing on a MinION ONT device (52). Despite showcasing great promise, the aforementioned strategies have not been applied to the more challenging setting of HIV-1 reservoir, which requires several orders of magnitude greater sensitivity.

We do acknowledge some limitations to this assay. First, the inclusion of the pre-amplification step to ensure efficient tagging and enrichment impedes the accurate quantification of reservoir clonality, as the dual UMI tags are only incorporated after the initial pre-amplification cycles. However, by performing multiple PCR replicates, we were still able to identify most clonal populations throughout this study. Further research into increasing the efficiency of the UMI tagging step would be needed to omit the pre-amplification step. Despite the limitations of HIV-PULSE in terms of accurate reservoir quantification, the assay can be valuable in areas where quantification does not matter, such as the aforementioned HIV-1 phylogenetics, drug resistance screening, and bNAb epitope mapping. Second, while we cannot exclude the possibility of chimera formation during the initial pre-amplification step, we consider it to be nearly impossible as chimera formation by PCR polymerase is normally observed in reactions with high numbers of PCR cycles (53). Third, we successfully applied the HIV-PULSE assay on samples from a chronic cohort of PLWH, yet samples from PLWH with different characteristics might be more challenging. As some steps of the analysis workflow rely on the sequence diversity to cluster identical bins into megabins to deconvolute the effect of pre-amplification, proviral reservoirs with lower intra-host sequence diversity (*e*.*g*. early ART initiation) could limit the success of this approach.

In conclusion, the HIV-PULSE assay presents itself as a promising HIV-1 NFL proviral sequencing method that enables scalable, high-throughput characterization of the proviral reservoir, while retaining sequencing accuracy comparable to HIV-1 NFL assays currently used in the field. We are convinced that the HIV-PULSE assay will be a valuable asset in advancing our understanding of the composition and dynamics of the viral reservoir during future basic and translational HIV-1 research.

## Data Availability

HIV-1 proviral sequences are uploaded to GenBank (accession numbers pending for GenBank approval). Sequencing data has been submitted to Sequence Read Archive (SRA) under BioProject ID (will be made available upon publication). The bioinformatics pipeline is available at https://github.com/laulambr/longread_umi_hiv.

## Acknowledgements

We would like to acknowledge and thank all participants who donated samples and all the clinicians and study nurses that assisted with the sample collection. We are grateful for the discussions with and input from Sarah Palmer, Bethany Horsburgh and Søren Karst. In addition, we would like to thank Ellen De Meester, Sarah De Keulenaer, and Sylvie Decraene from NXTGNT for their assistance in performing MiSeq sequencing. The following reagents were obtained through the NIH HIV Reagent Program, Division of AIDS, NIAID, NIH: J-Lat Full Length Cells (8.4), ARP-9847, contributed by Dr. Eric Verdin and Jurkat (E6-1) Cells, ARP-177, contributed by ATCC (Dr. Arthur Weiss).

## Table and Figure Legends

**Supplemental Table 1 Clinical characteristics of chronic cohort participants**.

**Supplemental Table 2 List of primers used throughout the study**.

**Supplemental Table 3 Performance results of the HIV-PULSE assay on participants of a chronic cohort**.

**Supplemental Table 4 Estimated costs per sequenced virus for FLIPS and HIV-PULSE**.

**Supplemental Figure 1 HIV-PULSE assay details and performance**.

(A) Visual presentation of the HIV-PULSE read construct layout. (B) Schematic representation of the bioinformatics workflow to analyze HIV-PULSE data. (C) Number of total reads for each HIV-PULSE sequencing run (LIB1 contained J-Lat 8.4 amplicon data, from LIB2 onwards clinical samples). (D) Sequencing library size (in base pairs) for each HIV-PULSE sequencing run. (E) Percentage of HIV-1 reads out of the total reads for each HIV-PULSE sequencing run. (F) Median read length of HIV-1 reads for each HIV-PULSE sequencing run. (G) Percentage of bins deemed correct out of the total detected bins for each HIV-PULSE sequencing run. (H) Percentage of reads belonging to correct bins out of the total number of binned reads for each HIV-PULSE sequencing run.

**Supplemental Figure 2 Proviral reservoir as assayed by FLIPS**.

(A) The proportions of different proviral classes observed among the distinct FLIPS proviruses for each participant. On the right the number of total and distinct proviruses, including proviruses that failed during *de novo* assembly is displayed for each participant (B) Distribution of the accuracy rates for all overlapping proviruses at different stages of the bioinformatics pipeline. The raw reads indicate the single-read accuracy (n= 232,131), racon3x and racon3x_medaka1x depict the HIV-PULSE bins (n=2,668) and megabins consists of the clustered HIV-PULSE bins (n=16). (C) Accuracy rates of overlapping proviruses detected with HIV-PULSE assay compared to their FLIPS Illumina reference counterpart. The color indicates the proviral genome classification by the HIV-PULSE assay for each respective provirus.

**Supplemental Figure 3 Bin coverage in function of amplicon length for all individuals**.

Each dot represents a single HIV-PULSE bin and is given a color based on the PCR-replicate. The dashed red line indicates the Q30 (99.9%) bin accuracy threshold.

**Supplemental Figure 4 Comparison of longitudinal sequencing data for P12**.

Phylogenetic tree of HIV-PULSE proviruses from P12 sampled at different timepoints (3 year interval). Tree was rooted against HXB2 and inversions were excluded (excluding 1 of the 21 timepoint overlapping clonal sequences). The symbols indicate the first (diamond) and second (circle) sampling timepoint while colors indicate whether the provirus was detected as clonal by the HIV-PULSE assay at that timepoint. The red arrows highlight identical proviral sequences detected at both timepoints.

**Supplemental Figure 5 Correlation between the number of distinct HIV-1 proviruses per PCR replicate and the total HIV-1 DNA reservoir size**.

For each participant, the number of distinct viruses for each sequenced PCR replicate are shown with the averages indicated as empty circles. A Spearman correlation (R=0.71, *p*=1.5*10^15^) is made between the mean number of distinct and total HIV-1 DNA copies/million CD4 cells as measured by ddPCR.

