## Supplemental Figures for "HIV-PULSE: A long-read sequencing assay for high-throughput near full-length HIV-1 proviral genome characterization"

### Supplemental Figure 1

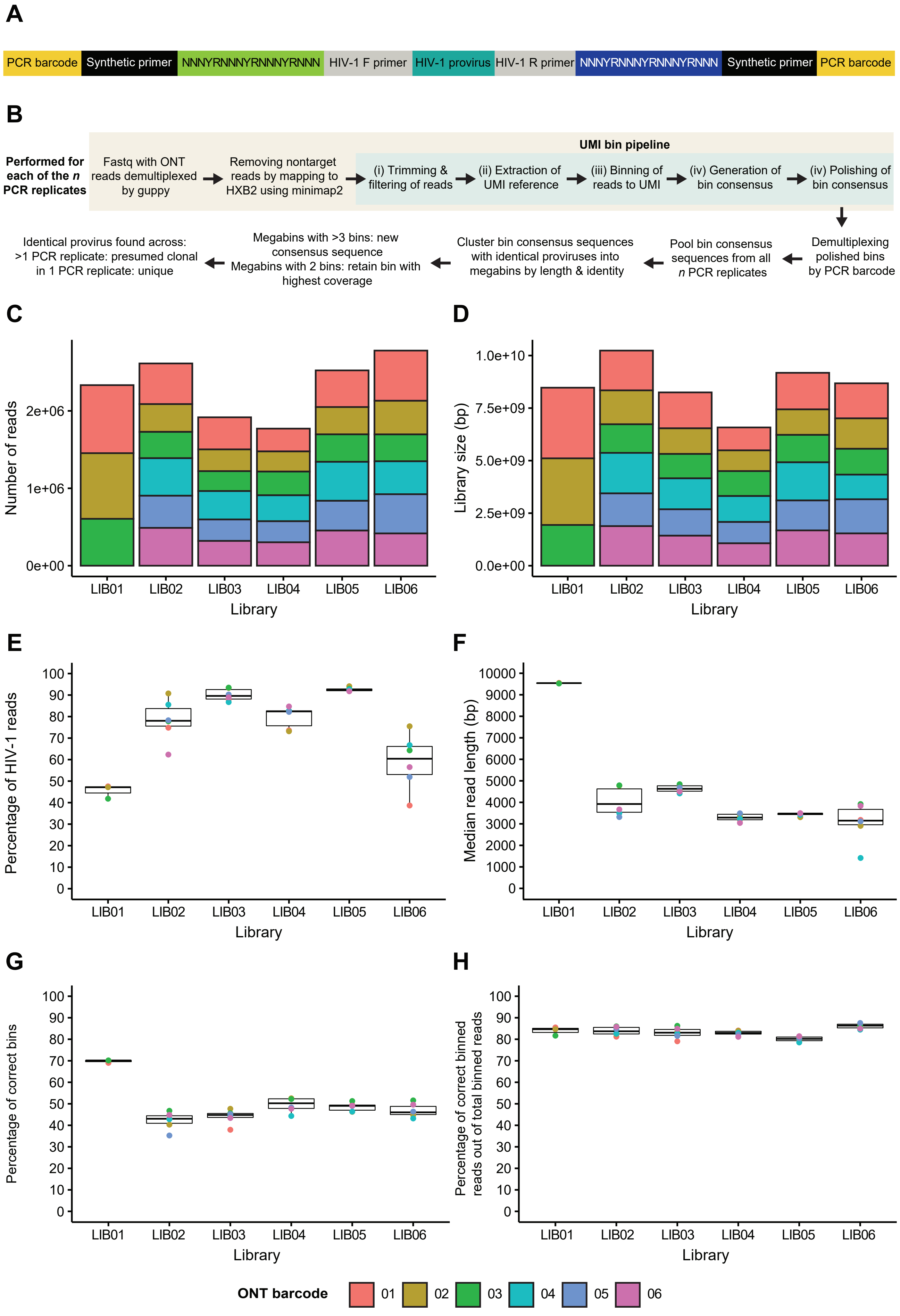

#### Supplemental Figure 2

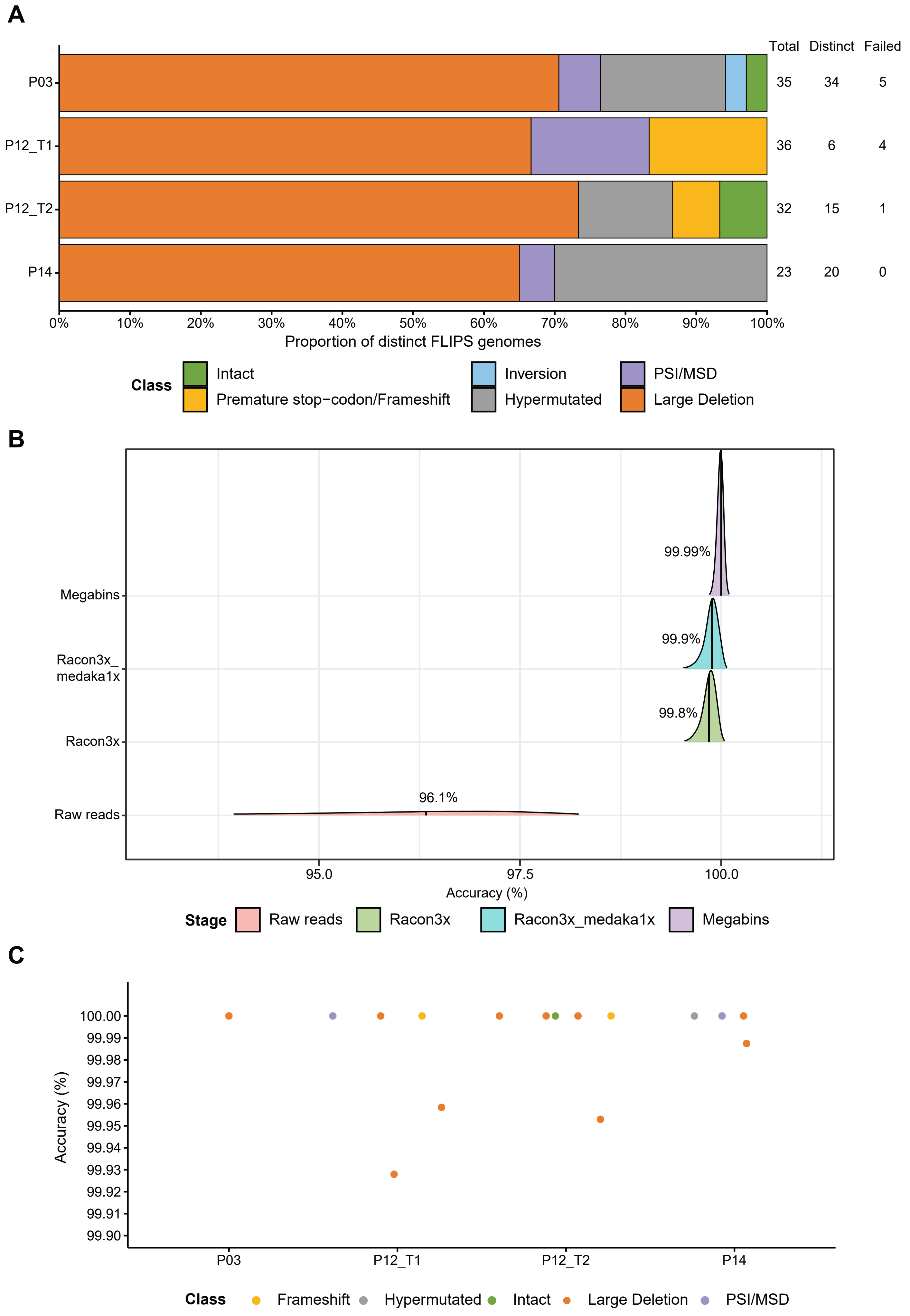

### Supplemental Figure 3

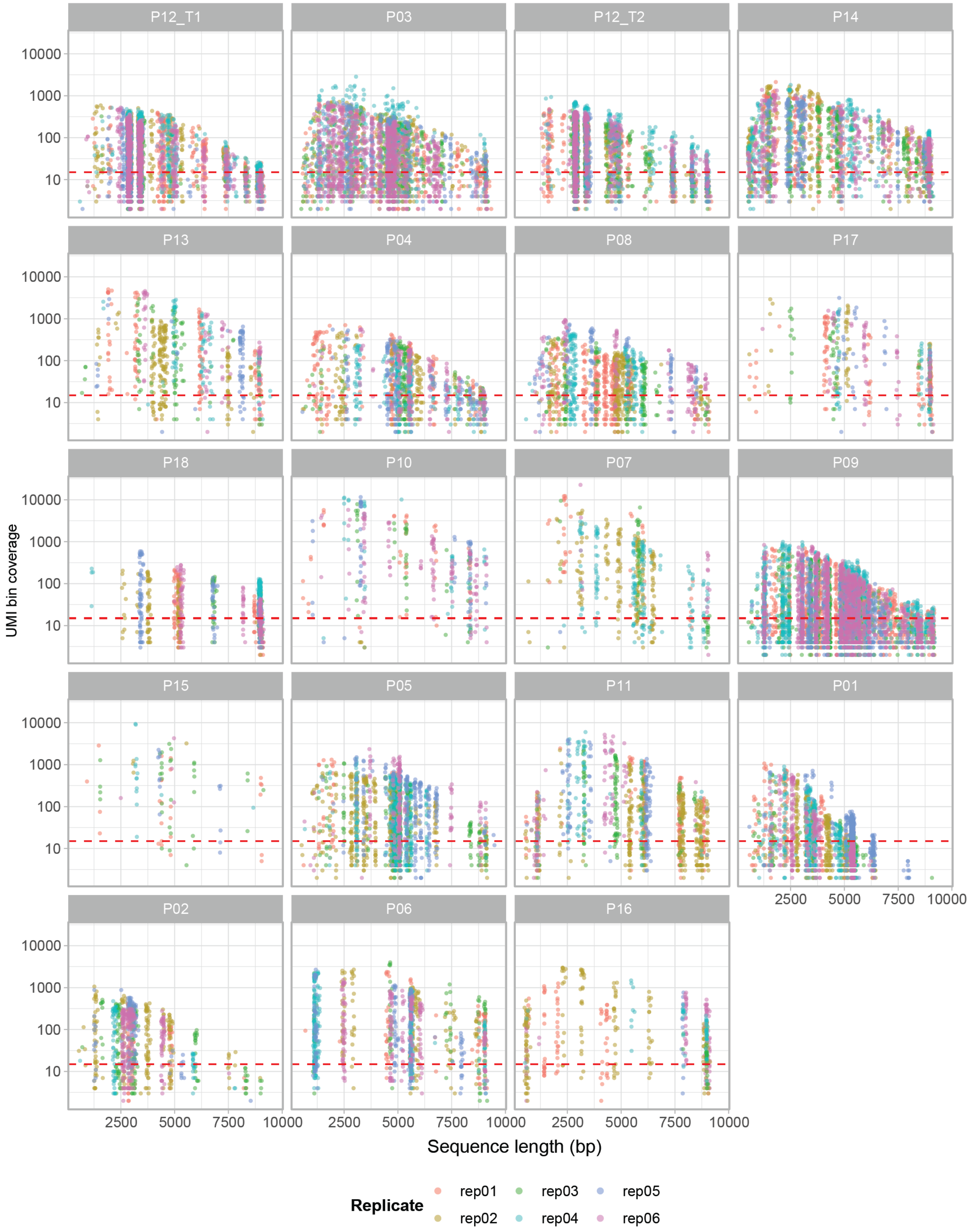

#### Supplemental Figure 4

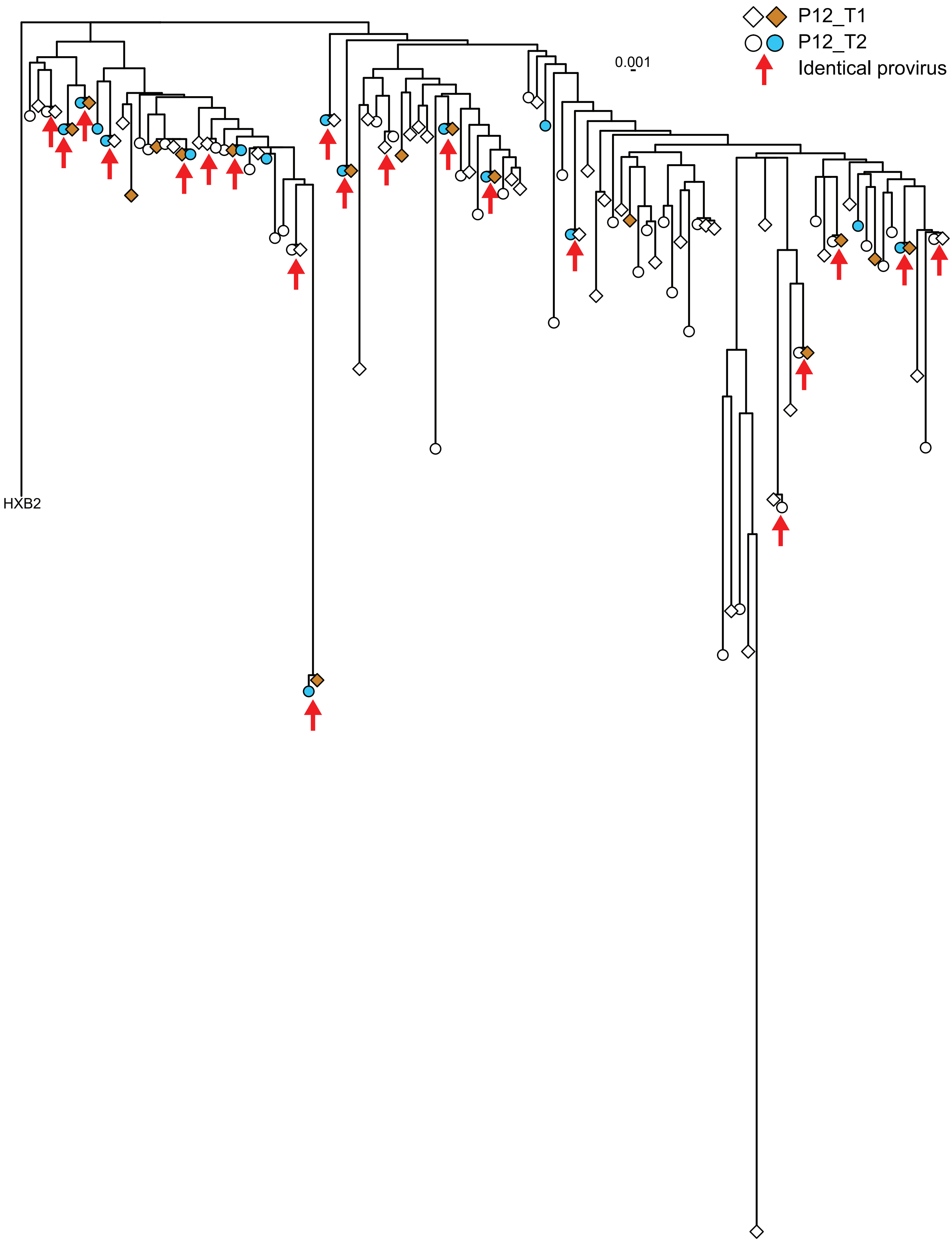

#### Supplemental Figure 5

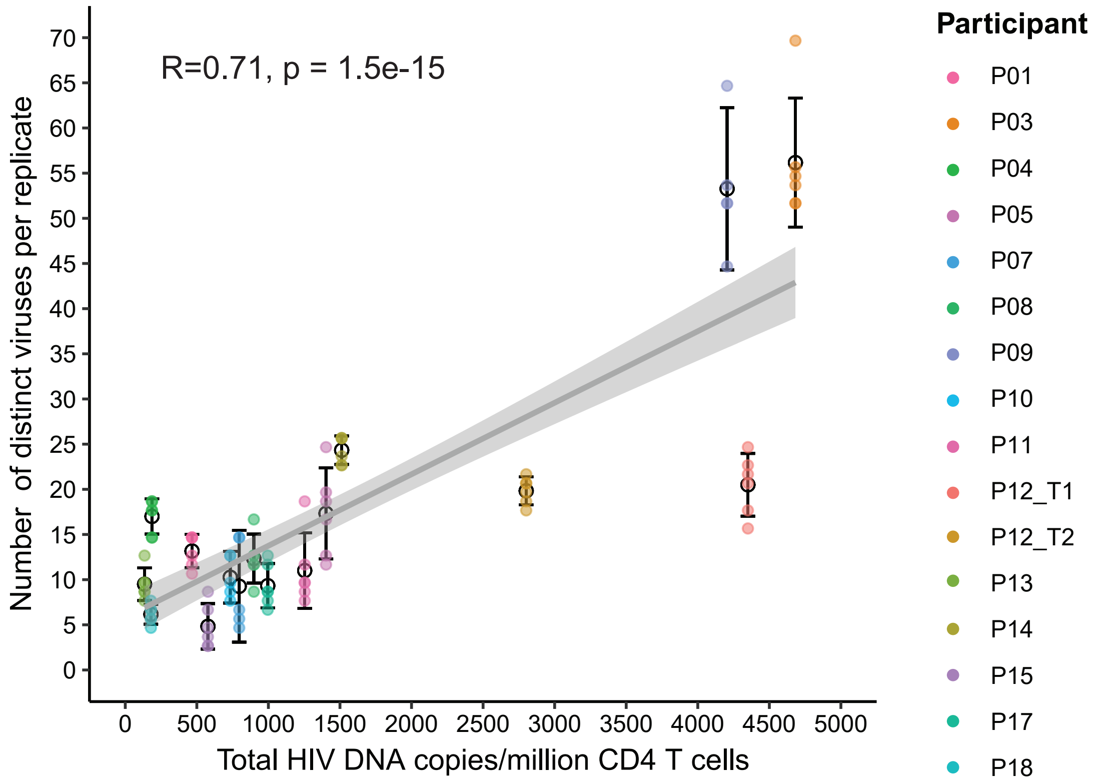
